# Extracellular vesicles from diverse fungal pathogens induce species-specific and endocytosis-dependent immunomodulation

**DOI:** 10.1101/2025.01.03.631181

**Authors:** Geneva N. Kwaku, Kirstine Nolling Jensen, Patricia Simaku, Daniel J. Floyd, Joseph W. Saelens, Christopher M. Reardon, Rebecca A. Ward, Kyle J. Basham, Olivia W. Hepworth, Tammy D. Vyas, Daniel Zamith-Miranda, Joshua D. Nosanchuk, Jatin M. Vyas, Hannah Brown Harding

## Abstract

Microbial pathogens generate extracellular vesicles (EVs) for intercellular communication and quorum sensing. Microbial EVs also induce inflammatory pathways within host innate immune cells. We previously demonstrated that EVs secreted by *Candida albicans* trigger type I interferon signaling in host cells specifically via the cGAS-STING innate immune signaling pathway. Here, we show that despite sharing similar properties of morphology and internal DNA content, the interactions between EVs and the innate immune system differ according to the parental fungal species. EVs secreted by *C. albicans*, *Saccharomyces cerevisiae, Cryptococcus neoformans,* and *Aspergillus fumigatus* are endocytosed at different rates by murine macrophages triggering varied cytokine responses, innate immune signaling, and subsequent immune cell recruitment. Notably, cell wall constituents that decorate *C. neoformans* and *A. fumigatus* EVs inhibit efficient internalization by macrophages and dampen innate immune activation. Our data uncover the transcriptional and functional consequences of the internalization of diverse fungal EVs by immune cells and reveal novel insights into the early innate immune response to distinct clinically significant fungal pathogens.

## INTRODUCTION

Invasive fungal infections have become more prevalent in recent years, largely impacting the growing population of immunocompromised individuals (1). Eukaryotic in nature, fungal infections are difficult to treat due to increasing antimicrobial resistance and the limited availability of antifungal therapeutics compared to their bacterial counterparts. The observed increase in mortality from fungal infection necessitates fundamental research on the molecular interactions between fungal pathogens and host organisms (2). Among others, the type I interferon (IFN) response has gained traction as a host defense against several common invasive fungal pathogens (3–5). In particular, the stimulator of IFN genes (STING) pathway is an orchestrated signaling cascade triggered by microbial pathogens that regulates IFN and interferon-stimulated genes (ISGs) during the early stages of infection. STING activation induces innate immune responses against several fungal organisms, including *Candida albicans* and *Aspergillus fumigatus* (6,7). Major IFN-producing pathways, including the STING pathway, are well studied in the context of viral and bacterial infections (8–13), however, their role during fungal infections remains understudied and represents a significant gap in knowledge. We recently identified the molecular mechanism behind *C. albicans* activation of the pathway. *C. albicans*-derived extracellular vesicles (EVs) trigger the induction of both IFNβ, a cytokine released upon STING pathway activation, and viperin, an ISG product, in a cGAS- and STING-dependent manner. *C. albicans* EVs and the DNA carried within them elicit an extensive proinflammatory response and phosphorylation of essential pathway components including IFN regulatory factor 3 (IRF3) and TANK-binding kinase1 (TBK1) (14). As whole fungal cells are typically sequestered into phagolysosomes upon host cell internalization, the fusion of *C. albicans* EVs (*Ca* EVs) to the plasma membrane provide a mechanism for how fungal nucleic acids access and activate the cytosolic DNA sensor cGAS. These findings led us to explore the interactions of additional species of fungal EVs with innate immune cells and their subsequent effects within host cells.

EVs are membrane-bound vesicles released from nearly all cell types, including both host and pathogen cells. They carry proteins, lipids, and nucleic acids and were first identified as messengers capable of shuttling information and cargo between the same cell types (15). It is now known that host cells can take up EVs released by pathogens and exploit the use of these microbial products to elicit robust and diverse pro-inflammatory or anti-inflammatory immune signaling in the host (16). In our study, we identified four different species of EV-generating fungi with diverse stimulatory, pathogenicity, and morphological properties to compare their early interactions with the innate immune system and any immunomodulatory impacts. Specifically, we examined the impact of cell wall constituents on EVs as virulence factors, selecting two species lacking significant cell wall constituents and two species possessing such components. *Saccharomyces cerevisiae* is a low-virulent yeast that generates EVs (*Sc* EVs) that do not contain robust cell wall constituents. *Sc* EVs have been studied in the context of cell wall remodeling and immune cell maturation (17,18). However, the mechanism of their internalization by macrophages and the extent to which *Sc* EVs trigger a type I IFN response is unknown (17,18). The encapsulated yeast *Cryptococcus neoformans* is an opportunistic fungal pathogen that can cause meningoencephalitis in immunocompromised individuals (19). Not only is this pathogenic yeast in a completely different phylum than *S. cerevisiae* and *C. albicans*, but it also possesses virulence factors and capsule that make it difficult to be phagocytosed. The presence of its surrounding capsule, comprised mainly of robust sugar molecules, glucuronoxylomannan (GXM), is a potent virulence factor of *C. neoformans* (20). GXM is present on *C. neoformans* EVs (*Cn* EVs), but the immunostimulatory properties of these EVs are understudied (21). Additionally, we explored fungal EV interactions with innate immune cells not only by yeasts, but also by a filamentous mold previously linked to type I IFN signaling, *A. fumigatus* (7). Although *A. fumigatus* conidia are not encapsulated by sugars like *C. neoformans*, they are surrounded by a hydrophobic layer of rodlet proteins, which may also be involved in immune evasion (22). Furthermore, *A. fumigatus*-derived EVs (*Af* EVs) prime murine macrophages for increased fungal clearance capacity and increased survival in the wax moth model *Galleria mellonella* after a fungal challenge (23,24). Thus, our study aimed to examine early host responses to EVs secreted from pathologically distinct fungal organisms that either possess or lack important cell wall constituents.

Our findings reveal novel patterns of macrophage ability to endocytose fungal EVs according to fungal species. *Ca* EVs and *Sc* EVs are more significantly endocytosed by macrophages compared to *Cn* EVs and *Af* EVs, the EVs that possess an outer polysaccharide capsule and a hydrophobic rodlet layer, respectively. In addition to our previous findings on *Ca* EVs, we identified EVs from diverse fungal organisms that differentially activate the STING pathway. The immunostimulatory properties of these EVs also differed by species, as macrophages stimulated by diverse EVs secreted different cytokines and differentially recruited neutrophils. Interestingly, the degree of pathway activation is EV species-dependent, revealing that EVs lacking structural outer layers are more efficiently endocytosed and can more robustly activate the STING pathway. As EVs may play a role in priming a host for infection and activating important proinflammatory mediators, these data provide new insights into the early host immune response to fungal pathogens and underscore the potential for fungal EVs to modulate innate immunity. By elucidating these interactions, this study generates the potential for new therapeutic targets and strategies in combatting fungal infections.

## RESULTS

### EV interactions with immune cells are species-specific

Since different species of fungi have unique impacts on host organisms, we inquired how interactions between fungal EVs and immune cells would compare between fungal species. We isolated EVs from *C. albicans*, *S. cerevisiae*, *C. neoformans,* and *A. fumigatus* cultures ***(Fig S1A)*** for co-culture with macrophages to compare EV internalization by innate immune cells. Interestingly, we observed the rate of EV uptake by macrophages to differ by species after a 3h co-incubation such that over 50% of the macrophage population internalized *Ca* EVs and *Sc* EVs, compared to less than 10% of macrophages having internalized *Cn* EVs and *Af* EVs (***Fig 1A***). To determine the mechanism of entry by the highly endocytosed *Ca* EVs and *Sc* EVs, we inhibited dynamin-dependent endocytosis in wildtype (WT) macrophages using dynasore (14,25). Dynasore treatment inhibited macrophages from endocytosing *Ca* EVs and *Sc* EVs by 80.4% and 88.9%, respectively, confirming endocytosis as the mechanism of EV entry into macrophages (***Fig 1B***).

**Fig 1.**
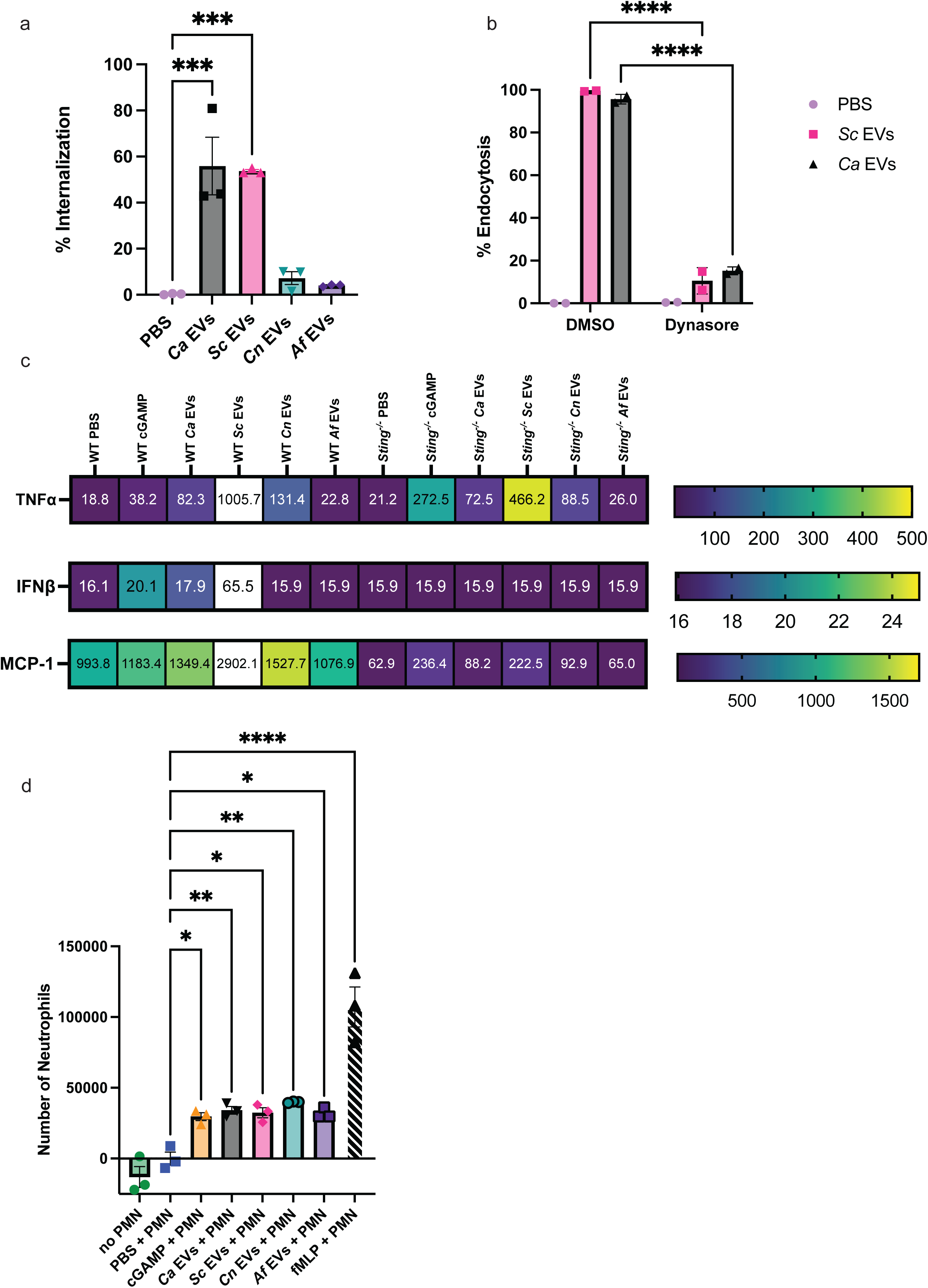
Murine macrophage internalization, cytokine production, and neutrophil recruitment differ in responses to different fungal EVs. **A,** Percent internalization of *Ca* EVs, *Sc* EVs, *Cn* EV*s*, and *Af* EVs by murine macrophages. Significance assessed by an ordinary one-way ANOVA and Dunnet’s multiple comparisons test, ***p=0.0002 vs PBS treated macrophages. n=3. **B,** Percent endocytosis of *Ca* EVs and *Sc* EVs with murine macrophages treated with DMSO or 100µM dynasore. Significance assessed by a two-way ANOVA and Tukey’s multiple comparisons test, ****p= <0.0001 vs respective DMSO controls. n=2. **C,** Heat maps of induction of TNFα, IFNβ, and MCP-1 (in pg/mL) in WT macrophages stimulated by PBS, 2.5µg cGAMP, *Ca*, *Sc, Cn,* and *Af* EVs. n=2. **D,** Number of neutrophils that transmigrated towards supernatants from WT macrophage stimulated by PBS, 2.5µg cGAMP, *Ca* EVs, *Sc* EVs*, Cn* EVs, and *Af* EVs (5 x 10^10^ EVs added per stimulation), and 0.1µM fMLP as a positive control. Significance assessed by an ordinary one-way ANOVA and Dunnet’s multiple comparisons test, *p≤0.0203, **p≤0.0073, ****p<0.0001 vs PBS-treated macrophages. n=3.

Given the differences in endocytosis of EVs by macrophages, we hypothesized that immune responses in innate immune cells would differ according to the species of fungal EV endocytosed. To address this hypothesis, we assessed the induction of 13 cytokines in macrophages exposed to fungal EVs. Since our previous findings revealed that *Ca* EVs activate the STING pathway in macrophages (14), we assessed levels of these 13 cytokines in both WT and *Sting^-/-^* macrophages stimulated with our fungal EVs, PBS, or cGAMP (positive control). Only three of the 13 cytokines showed changes compared to unstimulated macrophages (***Fig 1C* *and Fig S2A***): TNFα, IFNβ, and MCP-1. WT macrophages secreted higher levels of TNFα when stimulated by cGAMP, *Ca* EVs, *Sc* EVs, and *Cn* EVs (***Fig 1C, top panel***). Similarly to our prior results in mouse macrophages and human PBMCs (14), we observed an increased level of TNFα secretion in *Sting^-/-^* macrophages stimulated with *Ca* EVs compared to PBS-treated macrophages. This pattern was consistent with cGAMP, *Sc* EVs, and *Cn* EVs. We did not observe an increase in TNFα production in either *Af* EV-treated WT or *Sting^-/-^* macrophages. IFNβ was induced by cGAMP, *Ca* EVs, and *Sc* EVs in WT but not *Sting^-/-^* macrophages, or in macrophages stimulated with *Cn* or *Af* EVs (***Fig 1C, middle panel***). Increased MCP-1 secretion was induced by cGAMP and all EVs in WT, however, only cGAMP and *Sc* EVs induced moderate MCP-1 secretion in *Sting^-/-^* macrophages (***Fig 1C, bottom panel***).

Since MCP-1 plays a role in immune cell recruitment, we next assessed the recruitment of murine neutrophils in response to a co-culture of macrophage and fungal EVs. As a positive control for neutrophil recruitment, we used N-Formylmethionine-leucyl-phenylalanine (fMLP), a potent neutrophil chemoattractant (26). We revealed significant neutrophil recruitment in response to supernatants from macrophages co-cultured with all EVs, complementing MCP-1 induced by stimulation with the same EVs (***Fig 1D***). Interestingly, these data indicate that EVs derived from both yeast and filamentous fungi result in differential responses in macrophage host defense.

### Differential capacity of fungal EVs from diverse sources on STING pathway activation

Since we observed IFNβ to be highly induced by several fungal EVs and our previous work confirmed the induction of IFNβ in macrophages by *Ca* EVs as STING dependent (14), we next assessed STING pathway activation in macrophages stimulated by diverse pathogen-derived EVs. We assessed the expression of viperin, p-IRF3, IRF3, p-TBK1, and TBK1 as readouts of STING pathway activation. Viperin is an ISG product, and the phosphorylation of IRF3 and TBK1 is essential for the downstream transcriptional action of type I interferons and pro-inflammatory cytokines, which are key components of the host immune response. Like IFNβ production, we observed a significant increase in viperin induction in WT macrophages stimulated with *Ca* EVs and *Sc* EVs, with expression levels even greater than that from the cGAMP positive control **(*Fig 2A*).** Moderate viperin induction was observed in WT macrophages stimulated with *Cn* EVs, and minimal induction occurred in response to *Af* EVs (***Fig 2A***). As predicted, viperin expression in *Sting^-/-^*macrophages was absent upon EV stimulation, showing the necessity of this pathway for fungal EVs to induce viperin (***Fig S2B-C***). Furthermore, phosphorylation of the key downstream components of the STING pathway (IRF3 and TBK1) were induced to varying levels in response to fungal EVs in WT macrophages. The extent to which these proteins were phosphorylated was dependent on fungal species, exhibiting a similar expression pattern to viperin, where *Ca* EVs and *Sc* EVs induced greater phosphorylation compared to *Cn* EVs and *Af* EVs (***Fig 2A***).

**Fig 2.**
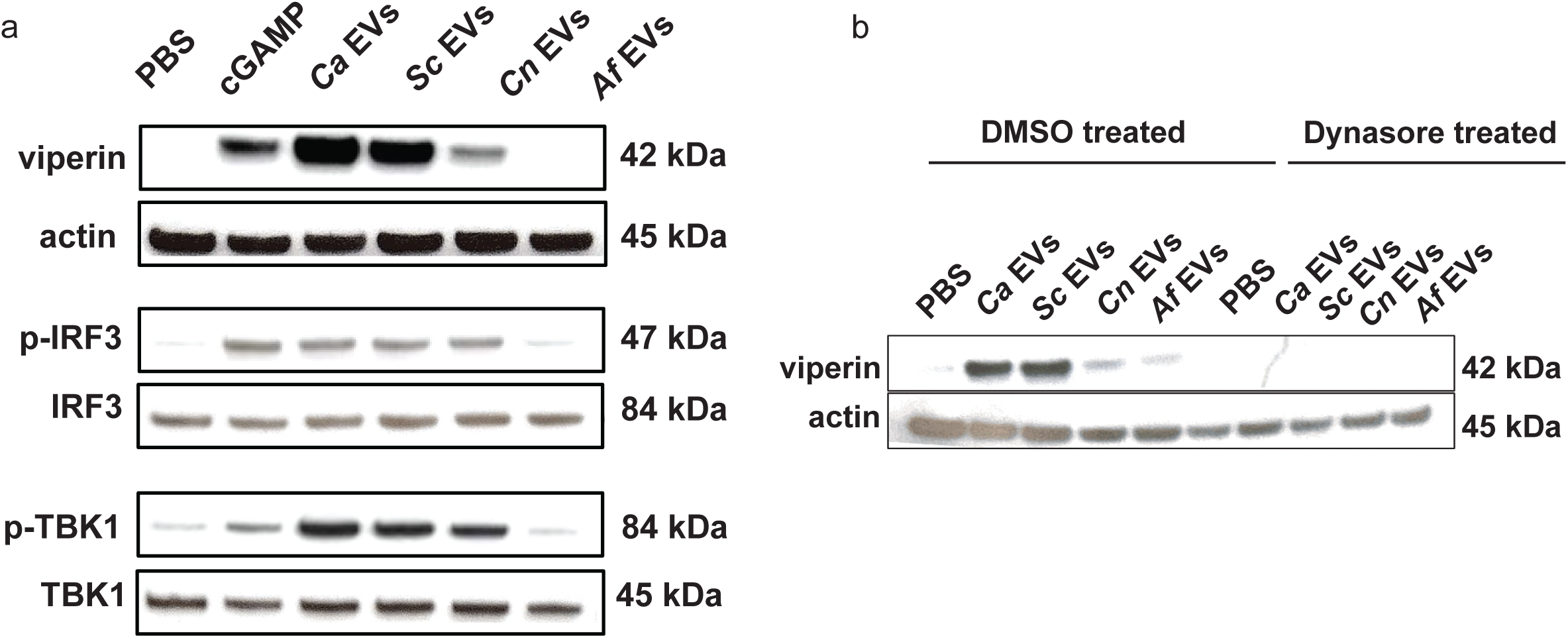
Fungal EVs differentially activate the STING pathway. **A**, Representative immunoblots of viperin, phosphorylated IRF3, total IRF3, phosphorylated TBK1, total TBK1, and actin in response to PBS, transfected cGAMP (2.5µg), *Ca* EVs, *Sc* EVs*, Cn* EVs, and *Af* EVs (5×10^10^ EVs/mL added per stimulation) in WT macrophages. **B**, Immunoblot of viperin in WT macrophages treated with either DMSO (solvent control) or 100µM dynasore and stimulated with *Ca* EVs, *Sc* EVs*, Cn* EVs, and *Af* EVs (5×10^10^ EVs/mL added per stimulation).

To assess whether this STING pathway activation relies on the endocytosis of EVs into macrophages, we next examined how preventing endocytosis would impact viperin production in macrophages stimulated with EVs using dynasore. Compared to the solvent controls which revealed a strong viperin induction in response to *Ca* EVs and *Sc* EVs, macrophages treated with dynasore showed no induction of viperin, regardless of stimulating EVs (***Fig 2B***). This further confirms the requirement of endocytosis for STING pathway activation and subsequent viperin induction by stimulating EVs.

### Translocalization of cGAS in response to fungal EVs

The localization of the DNA sensor cGAS is dynamic, as it remains tethered to the nuclear membrane in non-stimulatory conditions to prevent autoreactivity, and subsequently translocates to the cytosol upon stimulation in the presence of cytosolic DNA (27,28). Stimulation of macrophages expressing cGAS-GFP with *Ca* EVs increased non-nuclear localization of cGAS (14). Whether cGAS localization is impacted similarly across EVs from other fungal species remains unknown. Thus, we investigated how EVs from other fungal organisms impact cGAS localization. The lipid membranes of the EVs were fluorescently labeled with DiI, cultured with cGAS-GFP macrophages, and then imaged to assess the cGAS-localization in macrophages that had internalized EVs. Macrophages that endocytosed fungal EVs had greater non-nuclear localization of cGAS compared to the untreated macrophages without internalized EVs. However, the total percentage of non-nuclear cGAS localization varied among fungal species. The three yeast EVs (*i.e., Ca* EVs, *Sc* EVs, *Cn* EVs), induced noticeably more non-nuclear cGAS localization compared to *Af* EVs. Interestingly, *Sc* EVs induced the most non-nuclear localization of cGAS compared to all other EV types (***Fig 3A-B***). When cGAS was observed as non-nuclear, it was either localized solely cytoplasmic or a mix of cytoplasm and nucleus. While still inducing non-nuclear cGAS localization, the *Ca* EVs isolated from whole fungal organisms resulted in less localization of cGAS to the cytoplasm than we previously reported in stimulated macrophages with EVs isolated from *C. albicans* 48h grown biofilms (14). The mature nature of the 48h grown *C. albicans* biofilm EVs may explain the increased cGAS translocation, as a recent publication has noted changes in EV cargo content and membrane fluidics as a direct function of *C. albicans* biofilm growth time. Regardless, our results demonstrate the ability of fungal EVs to alter the cellular localization of the cGAS sensing molecule and these subcellular patterns reflect the ability of these EVs to induce downstream cGAS-STING signaling, viperin induction, and secretion of IFNβ.

**Fig 3.**
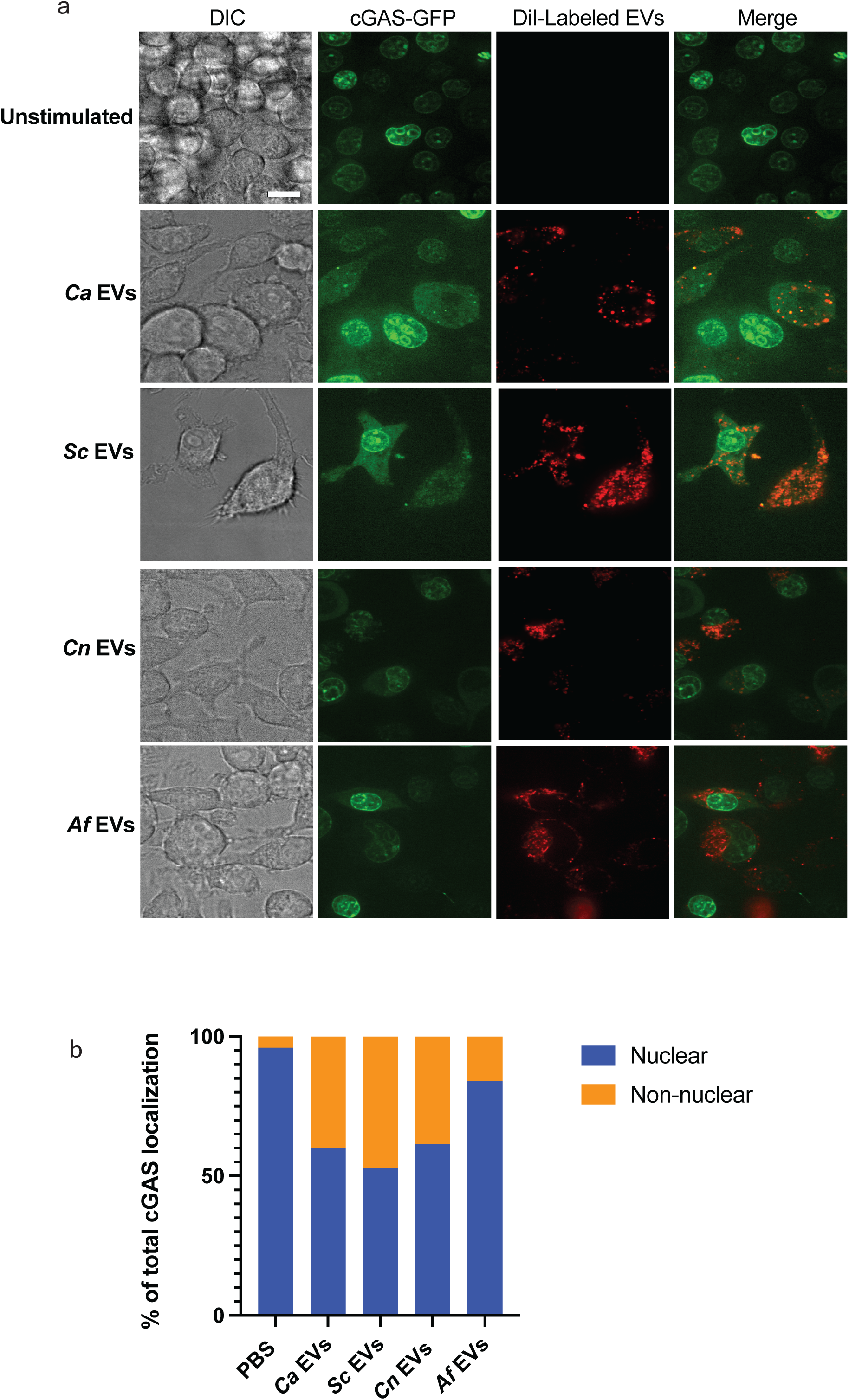
Fungal EVs induce translocation of cGAS from the nuclear membrane to the cytosol. **A,** Representative images of the localization of cGAS (green) when cGAS-GFP expressing macrophages are stimulated for 3h with DiI-labeled (red) PBS, *Ca* EVs, *Sc* EVs, *Cn* EVs, or *Af* EVs (8×10^9^ EVs added per stimulation). Scale bar=10µm. **B,** Semi-quantitative analysis of the percentage of cGAS localization that is either nuclear or non-nuclear. cGAS localization analyses were performed on approximately 100 cGAS-GFP expressing macrophages that successfully endocytosed DiI-labeled EVs.

### EVs share similar characteristics across diverse fungal species

EVs can vary in morphology and cargo depending on the source of the cell. Thus, to compare differences across our fungal species EVs, we analyzed several EV characteristics, including size, gross morphology, and due to the observed changes in STING pathway activation, internal DNA concentration and DNA sequences. Standard preparations of EVs from all species of interest yielded similar EV concentrations, typically between 1×10^10^ – 1×10^12^ EVs/mL. Although we noticed some variability, the differences were not statistically significant (***Fig 4A***). Interestingly, secreted EVs are known to be heterogenous in size, even when released by the same cell (29,30). EVs isolated from our fungal organisms ranged from 50-400 nm in diameter, with both the mode and median sizes of EVs from all four species falling within the 100-200 nm diameter range (***Fig 4B-C***). We next examined the gross morphology of our isolated EVs by transmission electron microscopy (TEM) to explore any significant structural differences between species (***Fig 4D***). All EVs appeared spherical with thick edges and noticeable depressions in the centers, a finding consistent with TEM imaging of EVs (31). There are no significant differences between species visible by TEM, indicating that fungal EV morphology is broadly conserved regardless of species or pathogenicity, including those that are historically non-pathogenic like *S. cerevisiae*, encapsulated like *C. neoformans*, and filamentous like *A. fumigatus*.

**Fig 4.**
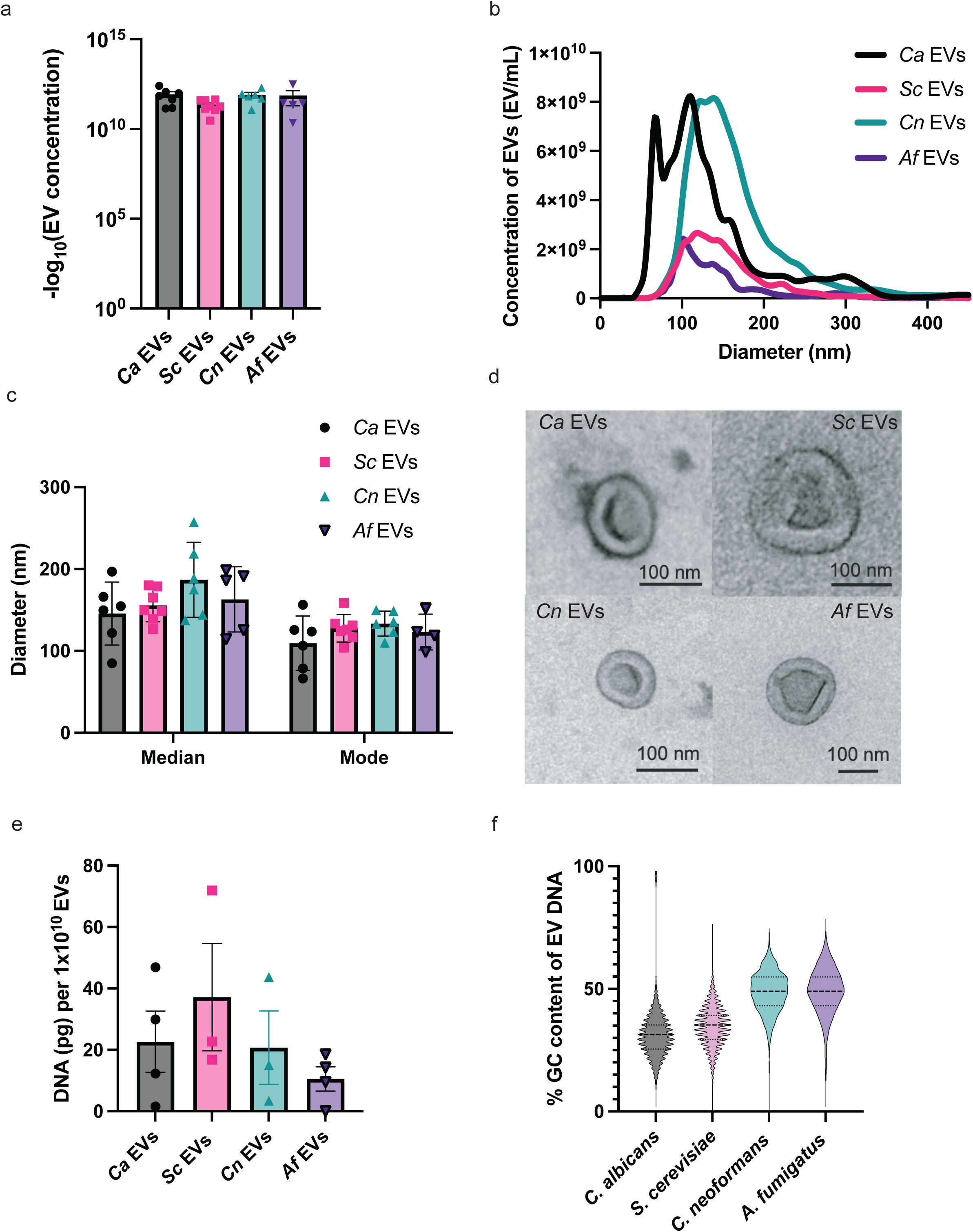
Physical and DNA characteristics of *Ca* EVs, *Sc* EVs, *Cn* EVs, and *Af* EVs. **A**, The - log_10_ values of EV concentration from standard isolation preps. Significance assessed using an ordinary one-way ANOVA and Tukey’s multiple comparisons test with no significance determined. n≥5. **B**, Quantification of EV concentration according to diameter for *Ca* EVs, *Sc* EVs, *Cn* EVs, and *Af* EVs. n≥5. **C,** The median and mode diameters of *Ca* EVs, *Sc* EVs, *Cn* EVs, and *Af* EVs. n≥5. **D**, Representative images from TEM of *Ca* EVs, *Sc* EVs, *Cn* EVs, and *Af* EVs. **E**, The average DNA concentration (pg per 1×10^10^ EVs) of *Ca* EVs, *Sc* EVs, *Cn* EVs, and *Af* EVs. Significance assessed using an ordinary one-way ANOVA and Tukey’s multiple comparisons test with no significance determined. n≥3. **F,** Percent GC content of EV DNA reads that mapped to the reference fungal genome as a truncated violin box plot.

We previously unveiled that DNA in *Ca* EVs triggers immune responses in macrophages through the STING pathway. With our findings that *Ca* EVs and *Sc* EVs are more significantly endocytosed by macrophages than *Cn* EVs and *Af* EVs, we hypothesized that EV DNA would be more abundant or stimulating in *Ca* EVs and *Sc* EVs compared to *Cn* EVs and *Af* EVs. Thus, we extracted DNA enclosed within the EVs, finding they contain similar concentrations across fungal species. The mean DNA concentration ranged from 10-40 pg per 1×10^10^ EVs (***Fig 4E***).

Finally, we more closely examined the DNA carried within these diverse EVs. Although there was no difference in quantity of DNA packaged in the different EVs (***Fig 4E***), we did note a difference in guanine-cytosine (GC) content between stimulatory and non-stimulatory EVs. We were interested in this aspect of EV DNA as GC-rich regions foster nucleosome formation, a structure that inhibits cGAS activation (32–38). Interestingly, DNA from *Ca* EVs and *Sc* EVs revealed lower compositions of GC content compared to *Cn* EV DNA and *Af* EV DNA ***(Fig 4F***). As *Cn* EVs and *Af* EVs very weakly triggered the DNA-dependent STING pathway, this finding is consistent with the hypothesis that nucleosome-favoring GC-rich DNA is less activating to cGAS, ultimately inducing reduced STING pathway activation. Therefore, not only could limited cGAS-STING pathway activation by *Cn* EVs and *Af* EVs be attributed to defects in endocytosis, but also to the increased GC content of the DNA carried within them.

### Outer structural layers on EVs impact endocytosis by macrophages

As we observed similarities in EV size and morphology from four different fungi, we concluded that other EV attributes may account for the observed differences in endocytosis rates by macrophages and subsequent degrees of STING pathway activation. One potential contributor may be physical differences in the outer EV membranes. To explore specific physical properties that might drive differences between EV internalization by immune cells and subsequent cGAS-STING pathway activation, we assessed the most external outer structural layers present on fungal EVs that had reduced internalization by macrophages, specifically the GXM polysaccharide capsule that surrounds *C. neoformans* yeast and the rodlet layer surrounding *A. fumigatus* conidia. We identified the presence of GXM on WT yeast cells and their EVs by confocal microscopy (***Fig 5A***) as previously described (39,40). We also used *cap59*Δ yeast, a *C. neoformans* strain that has significant reductions of the GXM layer (41,42). We confirmed the reduction of GXM on whole *cap59*Δ yeasts and their EVs by confocal microscopy (***Fig 5A***). To assess the effect of the rodlet layer on EV endocytosis, we used a mutant strain of *A. fumigatus* lacking this outermost layer, *ΔrodA*. This strain is known to yield conidia lacking the hydrophobic rodlet layer, however there is no rodlet antibody available to confirm via microscopy (43). We assessed the abilities of these mutant EVs to be endocytosed by macrophages by flow cytometry and found that those without outer cell wall constituents, *cap59*Δ *Cn* EVs and *ΔrodA Af* EVs exhibited significantly increased endocytosis compared to EVs with intact outer layers (***Fig 5B-C***). Furthermore, incubation of the *cap59*Δ EVs or *ΔrodA* EVs with WT macrophages yielded increased viperin expression compared to the respective WT EV counterparts (***Fig 5D-E***). Therefore, not only have we identified a species-specificity to the activation of the cGAS-STING pathway and downstream innate immune activation, but we also uncovered cell wall constituents as the suppressor agents preventing EV entry and robust STING pathway activation by *Cn* EVs and *Af* EVs. These findings further elucidate the mechanism by which EVs released by pathogenic fungi rewire the innate immune response.

**Fig 5.**
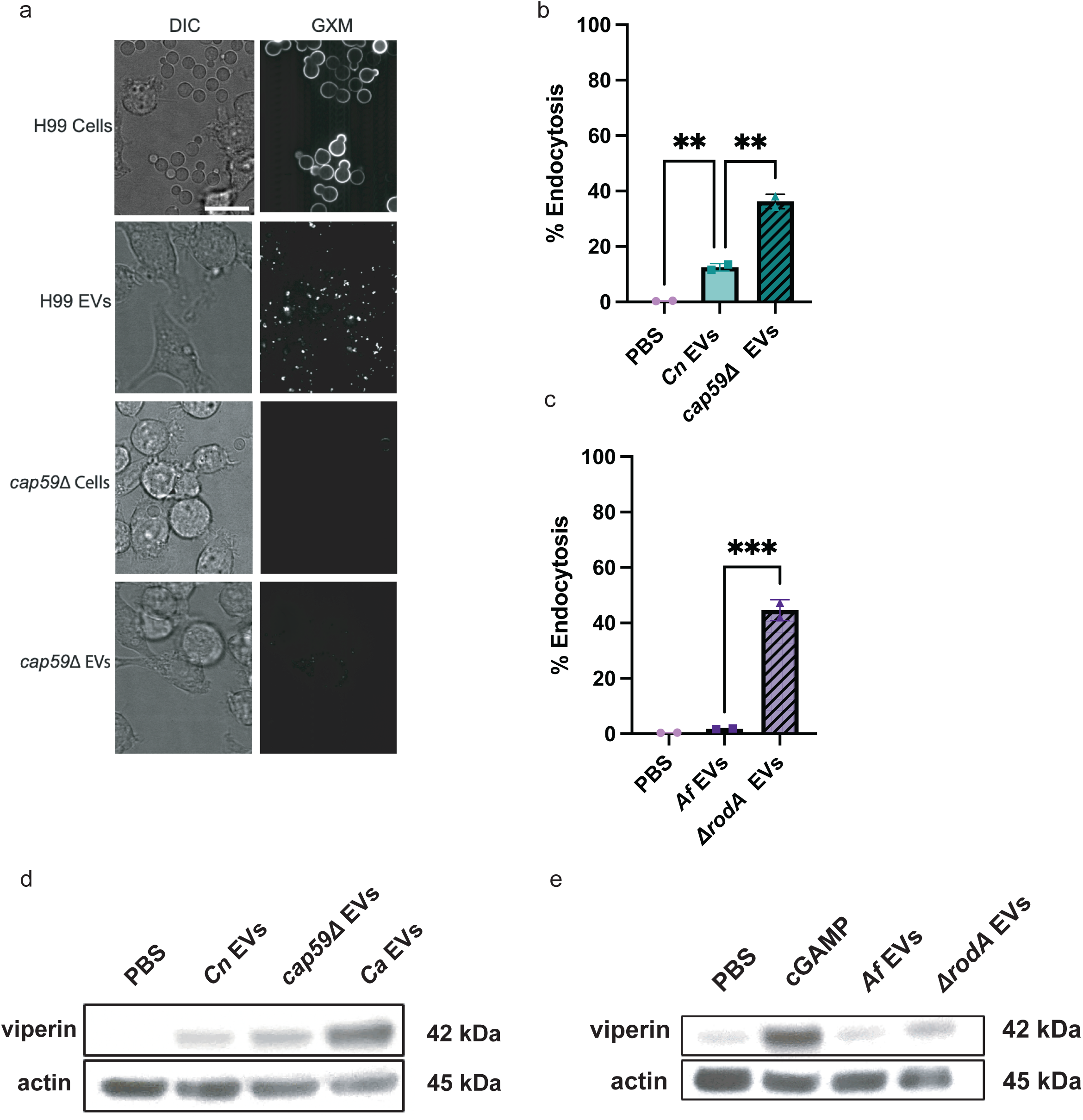
Outer structural layers on EVs inhibit endocytosis and STING pathway activation. **A,** Representative confocal microscopy showing GXM layer present on WT *C. neoformans* (H99) yeast and EVs but not on *cap59*Δ yeast and EVs. 1×10^7^ WT yeasts, 1×10^7^ *cap59*Δ yeasts, and 5×10^10^ WT and *cap59*Δ EVs were added to the respective stimulations. Scale bar=10µm. **B**, Percent endocytosis of *Cn* EVs and *cap59*Δ EVs by WT macrophages. n=2. **C,** Percent endocytosis of *Af* EVs and *ΔrodA* EVs by the total number of WT macrophages. n=2. **D,** Immunoblot of viperin and actin in WT macrophages stimulated by PBS, *Cn* EVs, *cap59*Δ EVs, and the positive control for yeast, *Ca* EVs (5×10^10^ EVs/mL were added per stimulation). **E,** Immunoblot of viperin and actin in WT macrophages stimulated by PBS, *Af* EVs, and *ΔrodA* EVs (5×10^10^ EVs/mL were added per stimulation), and positive control for *Aspergillus*: 2.5µg cGAMP.

## DISCUSSION

Here, we uncovered that the rate-limiting step for EV-mediated cGAS-STING pathway activation is endocytosis. EVs from *C. neoformans* and *A. fumigatus* are decorated with cell wall constituents that inhibit endocytosis, subsequently inhibiting cGAS-STING activation. While the immunomodulatory properties of cell wall constituents have been studied for whole fungal organisms (44–51), our findings provide novel insights connecting outer constituents and immune evasion in the context of fungal EVs. GXM and galactoxylolomannan (GalXM), major components of the outer capsule of *C. neoformans*, modulate both the innate and adaptive immune systems, including apoptosis induction of important inflammatory cells. For example, GXM and GalXM from *C. neoformans* induced macrophage apoptosis both *in vitro* and *in vivo,* a phenomenon mediated by Fas/FasL interactions (44). GalXM also interacts with glycoreceptors, triggering apoptosis of T cells via caspase-8 (45,46). Beyond influencing immune cell apoptosis, these capsular polysaccharides from various *Cryptococcus* species disrupt phagocyte migration, hinder phagocytosis, and inhibit NET production by human neutrophils, all contributing to immune evasion abilities of the encapsulated fungus (47–49). Similarly, surface proteins that create a hydrophobic layer over the surface of *Aspergillus* conidia are responsible for immune evasion tactics. WT *Aspergillus* conidia do not activate human dendritic cells and murine alveolar macrophages *in vitro*, whereas *ΔrodA* mutant conidia lacking the outer hydrophobic layer are highly stimulatory (50). The rodlet layer is also responsible for the dampening of pro-inflammatory signaling such as NF-κB and cytokine production via Dectin-1 and Dectin-2 (51). Additionally, in a murine infection model, mice infected with WT *A. fumigatus* resulted in a greater fungal burden compared to mice infected with *ΔrodA* (51).

Interestingly, another shared property of the outer layers of *Cryptococcus* and *Aspergillus* is the presence of melanin; *C. neoformans* produces melanin (3,4-dihydroxyphenylalanine melanin) from L-DOPA and *A. fumigatus* produces DHN melanin (1,8-dihydroxynapthalene melanin) (52–55). Notably, melanin increases microbial virulence and resistance to pharmacological treatment (56). Indeed, infection by *C. neoformans* lacking melanin-producing machinery prolonged host survival *in vivo* (*57,58*). Furthermore, melanin on *A. fumigatus* hinders CXCL1 and CXCL8 secretion and neutrophil recruitment from primary human airway epithelial cells by inhibiting calcium fluxing (59). In contrast to whole pathogens, much less is known about external structures surrounding microbial EVs. Interestingly, curvature-sensing peptides are used in EV detection procedures, where affinities of these peptides to bacterial EVs depend on the presence or absence of capsular polysaccharides on the EVs (60). Further studies elucidating extracellular structures of microbial EVs and how they impact immunity will be important for insight into host-pathogen relationships and tools for pathogen detection techniques.

Building on our previously published results demonstrating that dynamin-dependent endocytosis is necessary for the induction of viperin in macrophages exposed to *C. albicans* EVs (14), this study revealed a species-dependent EV endocytosis efficiency that directly correlates with activation of the cGAS-STING pathway. These observations are likely due to the protective and shielding nature of the cell wall constituents coating the outer layer of poorly stimulating fungal EVs. While this is the first time diverse fungal EV internalization by macrophages has been studied in the context of endocytosis and innate immune activation, the necessity for endocytosis of pathogens to elicit immune signaling has been studied extensively (61–63). Dynamin-dependent endocytosis is necessary for inflammatory cytokine (IFN*γ*, IL-6, and CCL5) induction in response to LPS (64). Dynamin-dependent endocytosis is also important for LPS-induced TLR4 internalization and initiation of downstream signaling (65). Additionally, dynasore-treated phagocytes have reduced induction of type III IFNs and decreased cell surface expression of CD86 and HLA class II molecules in *Streptococcus thermophilus*-stimulated cells (64). It is important to note that dynasore can work to disrupt dynamin-dependent endocytosis through inhibition of the GTPase activity of dynamin, blocking early constriction and fission, or by reducing plasma membrane cholesterol, which inhibits membrane mobilization (66). Therefore, it could be argued that the effects our study and others observed in dynasore-treated cells could be due to either altered early invagination or changes in macropinocytosis. Future studies into the specific endocytosis properties affected by our dynasore treated macrophages will elucidate the details of fungal EV internalization. However, these nuances do not diminish our overall finding that dynamin-dependent endocytosis is needed for efficient uptake of fungal EVs and downstream innate immune signaling and that this efficiency is affected by the outer layers of fungal EVs.

Importantly, initiation of cGAS-STING signaling is a critical priming step needed for the response to viral or foreign DNA (67). Recent evidence shows that cGAS binds to DNA via a “dual signal” model in which the subcellular localization of this interaction and the downstream immune activity initiated is dependent on the type of DNA (*i.e.,* foreign or self). Specifically, endocytosis of viral or other foreign DNA, an initial priming step occurs where spleen tyrosine kinase (SYK) and cGAS translocate to endosomes, where cGAS binds to the DNA and initiates a robust immune response. Alternatively, in response to the presence of self-DNA, cGAS localizes to the plasma membrane, which initiates a mild IFN response. Our studies did not investigate whether fungal EVs and EV DNA initiate this dual signal process and future experiments assessing the role of SYK in our model will elucidate whether this first priming step is essential following endocytosis of fungal EVs (68).

Finally, our studies further elucidate the differing abilities of fungal EVs to activate the STING pathway and type I IFN signaling. The STING pathway has historically been studied in the context of viral infections, bacterial infections, and autoimmunity (69). However, there are increasing reports of interactions between this pathway and fungal organisms. For example, *C. albicans* and *Ca* EVs activate the STING pathway and STING-deficient mice have improved survival following a *C. albicans* infection compared to their WT counterparts (14,70). *A. fumigatus* also elicits inflammatory responses in the host responsible for fungal keratitis via cGAS and STING (71). As EVs gain more attention as important mediators of pathogenic infection, it will be informative to explore the immunostimulatory capabilities of EVs alongside their whole organisms. Expanding our research questions to pathogenic fungi beyond those in our current study would contribute to a more comprehensive understanding of fungal properties that impact host-pathogen interactions. Our novel findings on the *in vitro* immunomodulatory abilities of *Ca* EVs, *Sc* EVs, *Cn* EVs, and *Af* EVs importantly inform early interactions with the innate immune system and provide details on infection mechanisms utilized by these clinically significant fungal pathogens.

## Supplemental Figures

**Fig S1.**
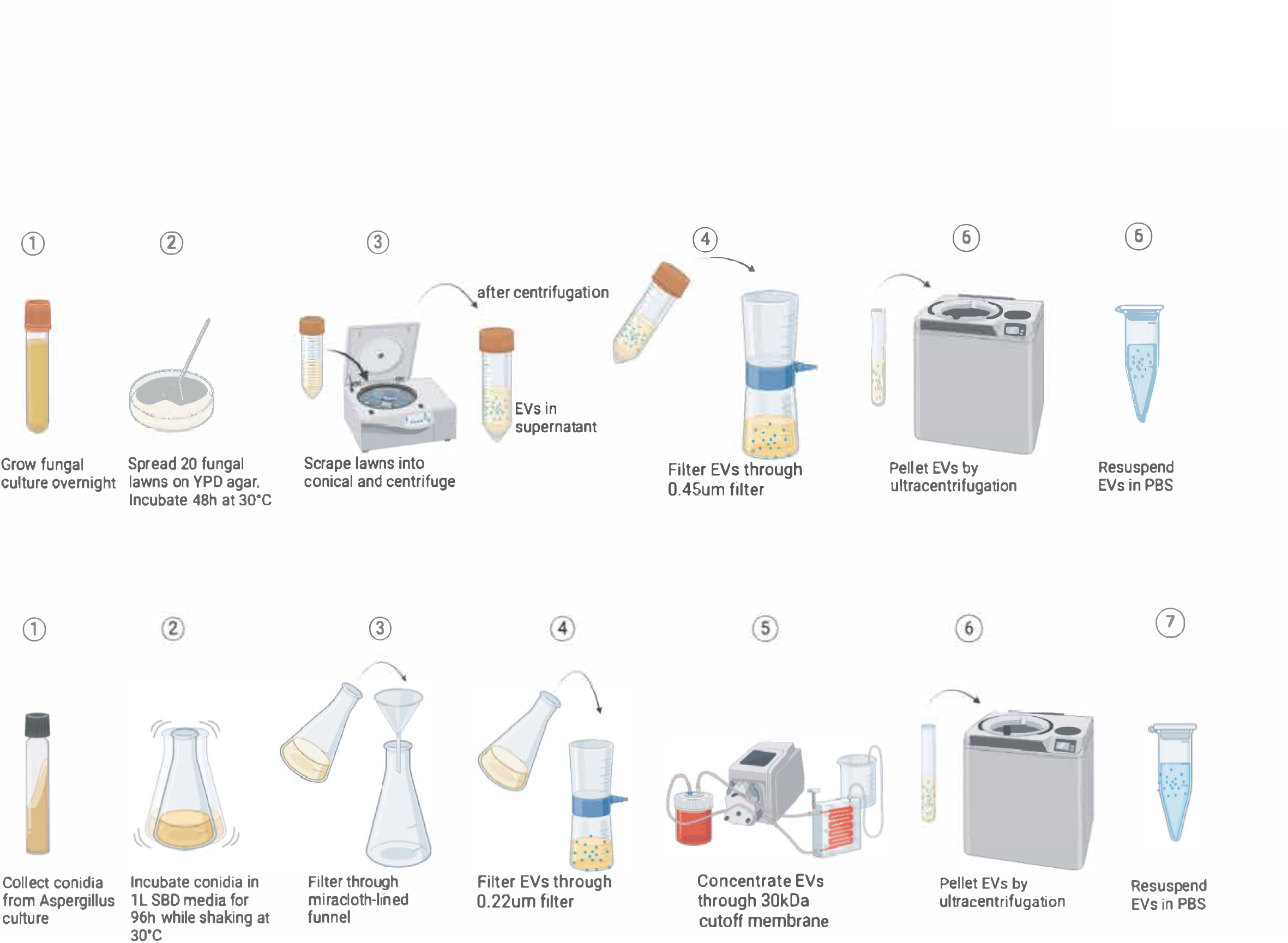
Isolation of fungal EVs. Isolation procedures of EVs from *C. albicans*, *S. cerevisiae*, *C. neoformans,* and *A. fumigatus* cultures.

**Fig S2.**
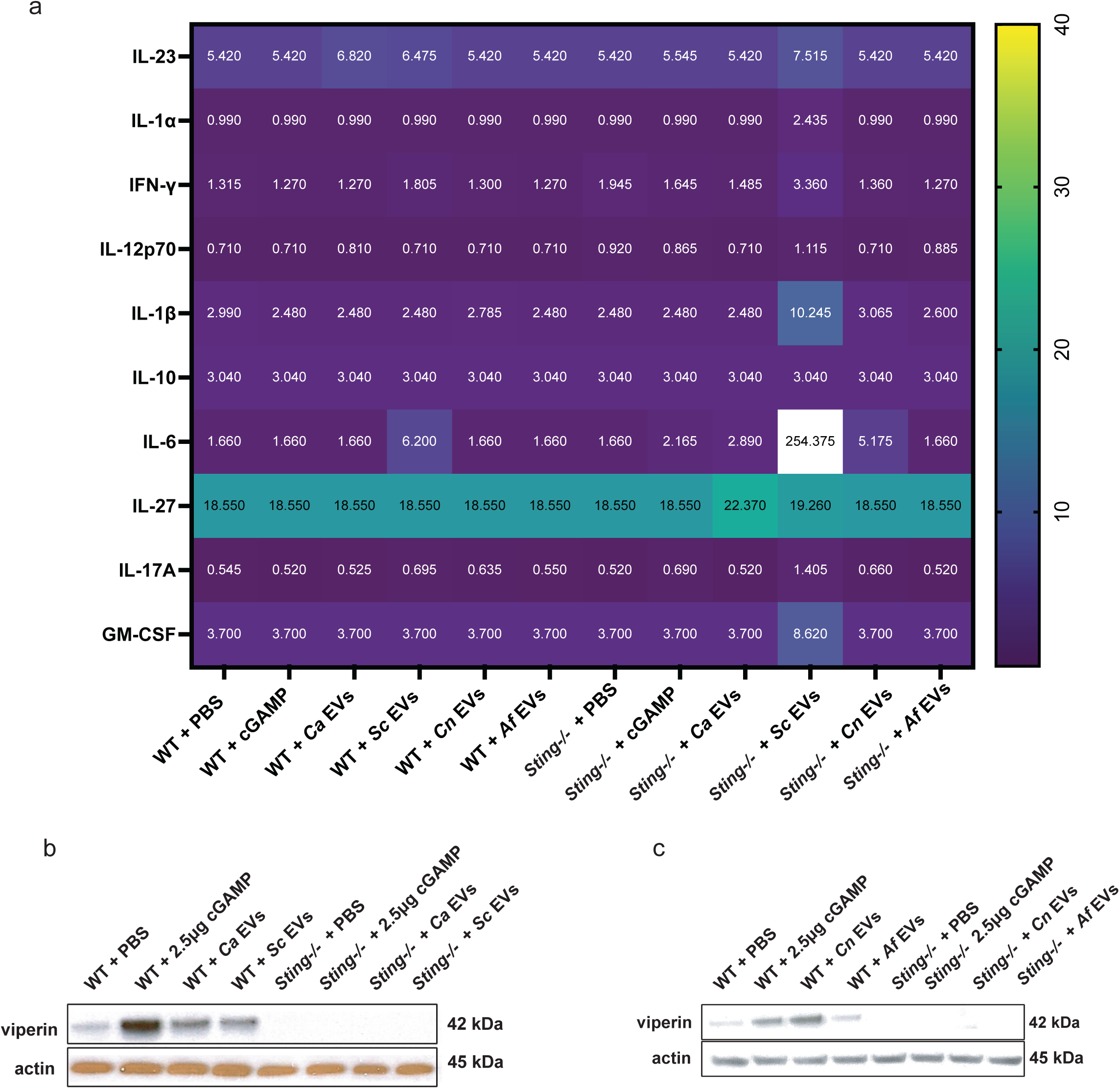
Immunomodulatory properties of fungal EVs. **A,** Heat map of selected cytokines secreted by WT and *Sting*^-/-^ macrophages when stimulated by PBS, cGAMP, *Ca* EVs, *Sc* EVs, *Cn* EVs, and *Af* EVs (EVs added at 1×10^10^ EVs/mL). **b,** Immunoblots of viperin and actin in WT and *Sting*^-/-^ macrophages stimulated by PBS, 2.5µg cGAMP, *Ca* EVs, or *Sc* EVs (EVs added at 1×10^10^ EVs/mL). **c,** Immunoblot of viperin and actin in WT and *Sting*^-/-^ macrophages stimulated by PBS, 2.5µg cGAMP, *Cn* EVs, or *Af* EVs (EVs added at 1×10^10^ EVs/mL).

## METHODS

### Cell culture and macrophage cell line generation

All macrophage lines were immortalized from C57BL/6 murine bone marrow cells. All immortalized bone marrow-derived macrophages (BMDM) were cultured in complete Dulbecco’s modified eagle medium (cDMEM) with added 10% FBS, penicillin and streptomycin (Pen+Strep), L-glutamine and 1M hydroxyethelpiperazine ethane sulfonic acid at 37°C and 5% CO_2_. Macrophages were washed with PBS and lifted by 0.05% trypsin in a 1:10 dilution. The WT immortalized BMDM were gifted by Douglas Golenbock (University of Massachusetts Medical School), and the *cGAS^-/-^* and *Sting^-/-^* macrophages were generated as previously described (72), as were the cGAS-GFP expressing macrophages.

### EV isolation and quantification

EVs from *C. albicans* (SC5314), *S. cerevisiae* (S228C), and *C. neoformans* (H99) were isolated and purified from YPD agar lawns. Each species was inoculated from a frozen stock in 5mL of YPD broth and incubated overnight at 30°C on a rotating wheel. Cells were washed in PBS, counted, and plated at 3.5×10^7^ cells/mL in 300µL per YPD agar plate, then spread by glass beads to form a lawn. A standard prep included 20 plates (STEMCELL Technologies 100mm x 15mm culture dish, non-treated) for each species. Plates were incubated at 30°C for 48h. The cell lawns were carefully scraped from the plates and transferred to conical tubes with 30mL of PBS. The conicals were centrifuged at 1,500*g* for 10 minutes to pellet the cellular debris. The EV-rich supernatant was filtered through 0.45µm filter. The filtered supernatants were collected (in 17mL Beckman Coulter centrifuge tubes Ref 337986) and ultracentrifuged at 100,000*g* for 1h at 4°C to pellet the EVs. The supernatants were aspirated, and the pellets were resuspended in 300µL each. EV sizes and concentrations were quantified using NanoSight LM10 & Nanoparticle Tracking Analysis (NTA) 3.4 analytical software (NanoSight Ltd., Minton Park, Amesbury, Wiltshire SP4 7RT, UK) and stored at −20°C for further use.

EVs from *A. fumigatus* (Af293) were isolated from liquid SBD broth. Af293 from a frozen stock was inoculated on GMM slants and incubated at 37°C for 72h and the conidia were harvested. Harvested conidia were inoculated at 2×10^8^ conidia/L into one liter of liquid SBD broth and incubated at 37°C for 96h at constant shaking at 150 rpm (New Brunswick Scientific I2500 Incubator Shaker). The culture was filtered through a miracloth-lined funnel and then through a 0.22µm filter overnight at 4°C. The filtered supernatant was concentrated by the Vivaflow 50 Concentrator with a 30kDa cut-off membrane. The concentrated supernatant was ultracentrifuged at 100,000*g* (in 17mL Beckman Coulter centrifuge tubes Ref 337986) for 1 hour at 4°C to pellet the EVs. The supernatants were aspirated and the pellets were resuspended in 300µL each. EV sizes and concentrations were quantified using NanoSight LM10 & Nanoparticle Tracking Analysis (NTA) 3.4 analytical software (NanoSight Ltd.,Minton Park, Amesbury, Wiltshire SP4 7RT, UK) and stored at −20°C for further use.

### Endocytosis analysis

WT immortalized macrophages were seeded at 1.5×10^5^ cells/mL in 1mL in a 12-well tissue culture treated plate and incubated overnight at 37°C and 5% CO_2_. We fluorescently labeled the lipid membranes of 5×10^10^ EVs from *C. albicans, S. cerevisiae, C. neoformans,* and *A. fumigatus* with DiI (DiI lipophilic stain (Invitrogen D282). 2μL of 1:100 dilutions of DiI in 100% ethanol were added to the EVs, followed by a 30 min incubation at RT. The EVs were then centrifuged at 100,000xG for 1h at 4°C (in 1.5mL Beckman Coulter microcentrifuge tubes REF357448) and then resuspended in 100μL to be co-incubated with the WT macrophages. A PBS-treated well was used for negative control. The stimulated macrophages were then incubated at 37°C and 5% CO_2_ for 3h. After incubation, we aspirated old media, washed with 1mL of PBS, aspirated and added 0.5mL of 0.05% trypsin to allow cells to detach from the well. We then centrifuged the samples to pellet the cells (1200rpm for 5 minutes), aspirated to remove the supernatant, and resuspended the pellet in 300μL PBS. Samples were filtered into FACS tubes to analyze the percent endocytosed EVs located in macrophages using a Cytek® Aurora Spectral flow cytometer (Cytek® Biosciences, California, USA). A total of 50,000 cells was collected per sample, recorded using the Spectroflo® (version 3.0.3, Cytek Biosciences). Finally, endocytosis was determined using negative unloaded macrophages and appropriate positive controls in duplicate. Data were analyzed using FlowJo™ (version 10.10.0, BD Biosciences, New Jersey, USA).

### Enzyme-linked immunosorbent assay (ELISA) analysis

To assess cytokine induction by macrophages co-cultured with EVs, we used a LEGENDplex™ 13-plex Mouse Inflammation Panel kit (cat. 740150). Macrophages were seeded at 1.5×10^5^ cells/mL in DMEM in a 12-well TC-treated plate overnight at 37°C and 5% CO_2_. Macrophages were stimulated with PBS, transfected cGAMP (2.5µg cGAMP), or pathogen-derived EVs at 1×10^10^ EVs/mL. Transfection for cGAMP was prepared with Lipofectamine 3000 reagents (Invitrogen L3000015). Supernatants from the co-incubations were collected after a 6h incubation at 37°C and 5% CO_2_, and the LEGENDplex™ Mouse Inflammation Panel protocol was followed accordingly to the kit instructions. The concentration of cytokines in each sample was determined using a BD FACSCelesta™ Flow Cytometer. This multiplex ELISA was completed in duplicate, and data were analyzed using the LEGENDplex™ Data Analysis Software Suite and PRISM10 software version 10.2.3 (GraphPad Software).

### Neutrophil Transmigration Assays

WT immortalized macrophages were seeded at 1.5×10^5^ cells/mL in 1mL in a 12-well tissue culture treated plate and incubated overnight at 37°C and 5% CO_2_. Macrophages were stimulated with 5×10^10^ EVs/mL or with cGAMP as a positive control. After 5 h of co-incubation at 37°C and 5% CO_2_, 500µL of the supernatant was transferred to the receiving plate of 24-well. N-Formylmethionine-leucyl-phenylalanine (fMLP) was added to one well of the receiving plate as a positive control. 3μm transwell inserts (Corning® Product No. 3415) were placed over the wells and 1 x 10^6^ murine neutrophils (immortalized cell line WT Cas9-ER-HoxB8-GMP), isolated as previously described (73), were added to the transwells in cRPMI media. Neutrophils were allowed to migrate for 2 h at 37°C and 5% CO_2_. The cells in the receiver plate were then lysed with Triton X while shaking for 20 min, and then treated with citrate buffer. 100µL of the samples were added in triplicate to a 96-well plate, and then treated with ABTS solution made from Di water, 2,2’-Azino-bis(3-ethylbenzo-thiazoline-6-sulfonic acid) diammonium salt (Sigma Aldrich) and citrate buffer. Neutrophil standards were also subjected to the same conditions. Absorbance was detected with an i3X spectrophotometer (Molecular Devices, LLC) at 405. Data were analyzed using PRISM10 software version 10.2.3 (GraphPad Software).

### Immunoblot analysis

For pathogen-derived EV and EV DNA stimulation experiments, immortalized macrophages were seeded at 1.5×10^5^ cells/mL in a 12-well or at 1.2×10^5^ cells/mL in a 48-well TC-treated plate overnight for cell adhesion at 37°C and 5% CO_2_. Pathogen-derived EVs were added to wells at 5×10^10^ EVs/mL or 1×10^10^ EVs/mL, and EV DNA was transfected at 300ng/mL. PBS was used as a negative control and cGAMP as a positive control. cGAMP and EV DNA were transfected using Lipofectamine 3000 reagents (Invitrogen L3000015). After a 6h incubation at 37°C and 5% CO_2_, lysates were collected by aspirating the media, washing cells with PBS, and then lysing cells with mammalian protein extraction reagent lysis buffer (Thermo Scientific, no. 78501) with sodium orthovanadate and protease inhibitors and shaking for 5 min. Lysates were collected and centrifuged at 14,000*g* for 5 min at 4°C. Lysates were transferred to new Eppendorf tubes with 4x NuPage lithium dodecyl sulfate loading buffer and 10x NuPage reducing agent. Protein expression was identified by western blot with 4-12% NuPage gel with 2-[N-morpholino]ethanesulfonic acid running buffer (NuPage gels, Thermo Fisher Scientific) at 180V for 1h, and transferred to methanol-activated polyvinylidene difluoride membrane (Perkin Elmer, Waltham, MA) with transfer buffer (0.025 M Tris, 0.192 M glycine and 20% methanol) and electrophoretic transfer at 100V for 1h. Membranes were blocked in 5% milk in 1X tris-buffered saline with Tween 20 (TBST) shaking for 1h at room temperature. For viperin detection, membranes were incubated rotating for 1 hour at room temperature in 1% BSA and a 1:5,000 dilution of primary antibody (Sigma MABF106) in 1X TBST. For TBK1 (Cell Signaling 3504) detection, membranes were incubated rotating for 1h at room temperature in 1% BSA and a 1:1,000 dilution of primary antibody in 1X TBST. For IRF3 (Cell Signaling 4302), membranes were incubated rotating for 1h at room temperature in 1% BSA and a 1:1,000 dilution of primary antibody in 1X TBST. For phospho-IRF3 (Cell Signaling E6F7Q) and phospho-TBK-1 (Cell Signaling D52C2) detection, membranes were incubated rotating overnight at 4°C in 5% BSA and 1:1,000 dilution of primary antibody in 1X TBST. After incubation of the primary antibody, the membranes were washed 3x with 1X TBST before incubation with secondary antibody swine anti-rabbit horseradish peroxidase-conjugated antibody at 1:2,000 (Agilent DAKO, P0399) or secondary peroxidase AffiniPure goat anti-mouse IgG (H and L) (Jackson ImmunoResearch) at 1:2,000 dilution in 1% milk in 1X TBST for 1h at room temperature. Membranes were then washed 3x with 1X TBST and then prepared with Western Lightning Plus ECL chemiluminescent substrate (Perkin Elmer) on Kodak BioMax XAR film (MilliporeSigma) for signal detection. Film sheets were scanned for electronic upload. Any contrast adjustments were applied evenly to the entire image and adhered to standards set forth by the scientific community. All reported Western blots/immunoblots were repeated in at least biological duplicate. Quantification of protein expression from western blots was performed with FIJI software version 1.53.

### EV DNA isolation

DNA was extracted from the isolated pathogen-derived EVs using the MasterPure Yeast DNA Purification Kit (Lucigen). 5 x 10^10^ EVs were added to 300 μL of yeast cell lysis solution and incubated at 65°C for 15 min. The samples were placed on ice for 5 min, and then 150 μL of milk protein concentrate protein precipitation reagent was added, vortex mixed for 10 seconds, and centrifuged in a table-top microcentrifuge for 10 min. The supernatants were transferred to new Eppendorf tubes, followed by the addition of 500 μL of isopropanol and mixed thoroughly by inversion. DNA was collected by centrifugation in a table-top microcentrifuge for 10 min at the maximum speed. The pellets containing the environmental DNA were washed with 500 μl of 70% ethanol, dried briefly at 42°C, and added to 50 μL of 1X TE buffer. DNA concentration was quantified by the Quant-iT PicoGreen dsDNA Assay Kit (Invitrogen) with fluorescent readings performed on an i3X spectrophotometer (Molecular Devices, LLC)

### EV DNA Sequencing

DNA from fungal EVs was extracted using Masterpure genomic extraction kit (above). DNA was submitted to the NextGen sequencing core at MGH and sequencing was done on a NextSeq Flowcell (P3 Single End 50 Run per million QC for Lanes after the First Lane). Reads that mapped to the reference genome were extracted with samtools view -h -F 4 and exported to fastq format with samtools bam2fq. The GC content analysis was done only the reads that mapped to the expected fungal reference genome. GC content was calculated from these reads by taking the sum of the counts of all G and C residues divided by the total number of bases.

### Confocal Microscopy

For the cGAS localization experiment, cGAS-GFP expressing macrophages were seeded at 1×10^5^ cells/mL in Nunc lab-tek II chambered coverglass (Thermo Scientific) in 500µL of cDMEM and incubated overnight at 37°C and 5% CO_2_. Macrophages were stimulated with fungal EVs at a concentration of 1.6×10^10^ cells/mL or stimulated with PBS. Before stimulation, 5μM DiI lipophilic stain (Invitrogen D282) was added to the EVs for 30 min at room temperature. The EVs were then ultracentrifuged at 100,000*g* for 1h at 4°C. A PBS sample also underwent the same staining and ultracentrifuge procedure. The macrophages were washed with PBS and refreshed with 500µL DMEM without FBS to rid of any serum-derived mammalian EVs. Finally, the macrophages were stimulated with the DiI labeled fungal EVs or the DiI-stained PBS and allowed to incubate for 3h at 37°C and 5% CO_2_. After incubation and phagocytosis of EVs, the cells were imaged on a Nikon Inverted Microscope Eclipse Ti-E with a CSU-X1 confocal spinning disk head (Yokogawa), and a 6-laser Cairn Multiline launch equipped with 405nm, 445nm, 488nm, 515nm, 561nm, and 647nm solid-state lasers was used to excite the sample. The objective was a 100x high-numerical aperture objective (Nikon, 1003, 1.49 numerical aperture, oil immersion) and an EMCCD camera (Hamamatsu Photonics). For image acquisition, we used MetaMorph software version 7.10.5.476 (Molecular Devices). The photos were cropped in FIJI software and organized in Adobe Illustrator, version 28.5 (Adobe Systems). The brightness/contrast values were set to the recommended “auto” levels for each individual image. The localization of cGAS in the macrophages was quantified as “nuclear” if cGAS was only present in the nucleus, “cytosolic” if cGAS was only present in the cytoplasm, or “mixed” if cGAS was only present in both the nucleus and cytoplasm.

For the GXM staining, WT macrophages were seeded at 1×10^5^ cell/mL in 500µL of cDMEM in Nunc lab-tek II chambered coverglass (Thermo Scientific) and incubated overnight at 37°C and 5% CO_2_. 10uL of antiGXM antibody (Millipore Sigma Cat. No. MABF2069) was added to 1×10^7^ H99 cells and 1×10^6^ cap59Δ yeast cells. 10uL of antiGXM antibody (Millipore Sigma Cat. No. MABF2069) was also added to 5 ×10^10^ H99 EVs and 5 ×10^10^ cap59Δ EVs, followed by a 1-hour incubation of all four of these samples at 37°C while shaking. 5uL of secondary anti-mouse AF488 (ThermoFisher Scientific Ref A32723 Lot WF323912) antibody was then added to the samples, followed by another 1-hour incubation at 37°C while shaking. Samples were ultracentrifuged for 1 hour at 4°C at 100,000g. Supernatant was removed and the pellets were resuspended in 50uL of PBS which was then added to the plated macrophages and co-incubated for 3 hours. Images were then captured by confocal microscopy and analyzed as previously described.

### TEM imaging (Negative Staining)

10-20µL of EV samples at approximately 1×10^11^ EVs/mL were added onto 200 mesh Formvar/carbon coated nickel grids and allowed to adsorb 15min. Grids were blotted to remove excess suspension, rinsed briefly with filtered distilled deionized water, and contrast-stained for 10min in a tylose/uranyl acetate solution on ice. Grids were blotted again and allowed to air dry prior to analysis. Examination of preparations was done using a JEOL JEM 1011 transmission electron microscope at 80kV. Images were collected using an AMT digital imaging system with proprietary image capture software (Advanced Microscopy Techniques, Danvers, Massachusetts).

## ACKNOWLEDGMENTS

The work in the Vyas Lab is supported by grants from the NIH: R01AI150181, R01AI136529, R21AI152499 (JMV). JN and DZM are supported by R01AI171093 and R21AI156104. Electron microscopy was performed in the Massachusetts General Hospital (MGH) Microscopy Core of the Program in Membrane Biology, which is partially supported by an Inflammatory Bowel Disease Grant DK043351 and a Boston Area Diabetes and Endocrinology Research Center (BADERC) Award DK135043. DNA Sequencing was performed by the NextGen Sequencing Core at MGH. Nanoparticle tracking analysis was performed at the Nanosight Nanoparticle Sizing and Quantification Facility in the Microscopy Core of MGH. The authors would like to thank all members of the Mansour Laboratory (MGH) for engaging discussions and Marta Brandt (Broad Institute of MIT and Harvard) and Ramnik J. Xavier (Broad Institute of MIT and Harvard & MGH) for assistance with the Cytek® Aurora Spectral flow cytometer.

## Notes

### Competing Interest Statement

The authors have declared no competing interest.

